# A Deep Hypergraph Learning Model for Predicting Antimicrobial Combination Effects Across Bacterial Targets

**DOI:** 10.64898/2026.06.09.731104

**Authors:** Farzad Midjani, Amir Hossein Rajabi, Fateme Zahra Keshtkar, Mahdi Malekpour, Ali Jafarizadeh, Roohallah Alizadehsani, Pawel Plawiak

## Abstract

Antimicrobial resistance (AMR) creates an urgent need for efficient strategies to identify effective antibacterial combinations. Combination therapy, including antimicrobial peptides (AMPs) paired with conventional antibiotics, is a promising approach, but exhaustive experimental screening across drug pairs and bacterial targets is impractical. This study introduces a hybrid GCN-based hypergraph neural network (HGNN) for predicting antimicrobial-agent combination outcomes against bacterial targets. Each antimicrobial-agent-antimicrobial-agent-bacterium triplet is represented as a ternary hyperedge, enabling the model to learn context-dependent interaction patterns.

The framework integrates SMILES-derived molecular graph embeddings for antimicrobial agents, including conventional antibiotics and AMPs, with taxonomy-derived bacterial representations. The prediction task was formulated as a three-class classification problem: synergy, antagonism, and non-interaction. The non-interaction class included experimentally verified indifferent records and synthetic presumed non-interaction triplets generated by negative sampling. Model development used drug-pair-grouped splitting, five-fold grouped cross-validation within the training/validation partition, and final evaluation on a held-out test set.

On the held-out three-class test set, the selected GCN-based HGNN achieved an accuracy of 0.83, weighted F1-score of 0.84, macro F1-score of 0.80, and ROC-AUC of 0.95. Per-class evaluation showed accuracies of 0.80 for synergy, 0.92 for antagonism, and 0.85 for non-interaction. Pair-type analysis showed strong performance across AMP-AMP, AMP-conventional antibiotic, and conventional antibiotic-conventional antibiotic combinations.

These findings suggest that hypergraph-based representation learning can support computational prioritization of antimicrobial combinations for experimental follow-up. Further studies will be needed to improve model interpretability and to perform prospective validation of predicted synergistic combinations.

## 1 Introduction

Antimicrobial resistance (AMR) is a major global health threat that is reducing the effectiveness of established antibacterial treatments [1]. Drug-resistant bacterial infections were associated with an estimated 4.95 million deaths in 2019, and future projections suggest that the burden could increase substantially without coordinated intervention [2–4]. Overuse and misuse of antimicrobial agents in human medicine, veterinary practice, and agriculture, together with international travel and trade, contribute to the emergence and spread of resistant bacterial strains [1, 5–9].

Bacterial resistance arises through mechanisms such as enzymatic drug inactivation, target modification, reduced intracellular accumulation, active efflux, and horizontal transfer of resistance genes [10–13]. These mechanisms contribute to multidrug-resistant, extensively drug-resistant, and pandrug-resistant infections, which limit treatment options and are associated with poorer clinical and economic outcomes [3, 14].

The effects of drug combinations are typically categorized as synergistic, additive, or antagonistic [15]. Identifying synergistic combinations is important, as these offer the greatest potential for therapeutic benefit [16]. However, the experimental screening of all possible drug combinations against a multitude of bacterial pathogens is impractical at scale: given the vast combinatorial space, testing every pair (or higher-order combination) against hundreds of clinically relevant bacterial strains and their resistant variants rapidly becomes economically and logistically infeasible using traditional laboratory methods such as checkerboard assays or time-kill studies [17, 18].

AMPs are naturally occurring components of the innate immune system in virtually all forms of life. Their synergistic effects, for example, permeabilizing bacterial membranes to facilitate the entry of other drugs or targeting cellular pathways distinct from those affected by conventional antibiotics, make them an attractive choice for combination therapy [19,20]. However, predicting AMP-containing combinations, particularly AMP-conventional antibiotic interactions, presents additional challenges for computational models. AMPs are structurally diverse, ranging from small linear peptides to more complex cyclic or disulfide-bonded structures [19]. Moreover, data on AMP combinations, while growing, are still less extensive than for conventional antibiotic combinations [20, 21].

Over the years, different methods have been applied to this issue. Some earlier approaches relied on quantitative structure-activity relationship (QSAR) models to correlate the physicochemical properties of drugs with their interaction outcomes [22]. Another strategy involved using pharmacokinetic/pharmacodynamic (PK/PD) models to simulate drug exposure and effects over time [23]. Recently, machine learning models have been applied, using features derived from drug structures, targets, mechanisms of action, gene expression profiles, or other biological data to train predictive models [24, 25]. Despite their promise, these approaches simplify complex biological interactions and may not fully capture the nuances of combination effects. Standard GNNs are inherently designed to model pairwise relationships [26]. When applied to drug combinations, this typically involves predicting an interaction between two drugs, or a drug and a target [27]. However, an antimicrobial combination effect (e.g., synergy between Drug A and Drug B against Bacterium X) is fundamentally a three-way interaction [28–30]. The outcome depends not just on Drug A and Drug B, but critically on the specific bacterium they are acting against [28–30]. Forcing this three-way relationship into a pairwise framework can lead to loss of information and an inability to capture the context-dependent nature of these higher order interactions [31]. This motivates modeling triplets explicitly rather than via pairwise decompositions.

Hypergraph neural networks (HGNNs) naturally represent higher-order relations by allowing each hyperedge to connect more than two nodes [32]. In antimicrobial combination prediction, this allows each antimicrobial agent-antimicrobial agent-bacterium triplet to be encoded as a ternary hyperedge, directly representing the joint interaction between two antimicrobial agents and a bacterial target. Unlike conventional pairwise graphs, this structure enables the model to learn context-dependent representations influenced by all three components [31, 33, 34].

Here, we propose a GCN-based hypergraph framework that integrates SMILES-derived molecular embeddings for antimicrobial agents with taxonomy-derived representations for bacterial targets. The model predicts three interaction classes: synergy, antagonism, and non-interaction. By preserving the ternary hyperedge structure, the framework captures antimicrobial combination effects as outcomes that depend jointly on both agents and the bacterial target.

## 2 Methods

The workflow consisted of data acquisition and preprocessing, feature construction for antimicrobial agents and bacterial targets, hypergraph formulation, model training, and held-out evaluation, as summarized in Figure 1. The prediction task was formulated as a three-class classification problem: synergy, antagonism, and non-interaction.

**Figure 1.**
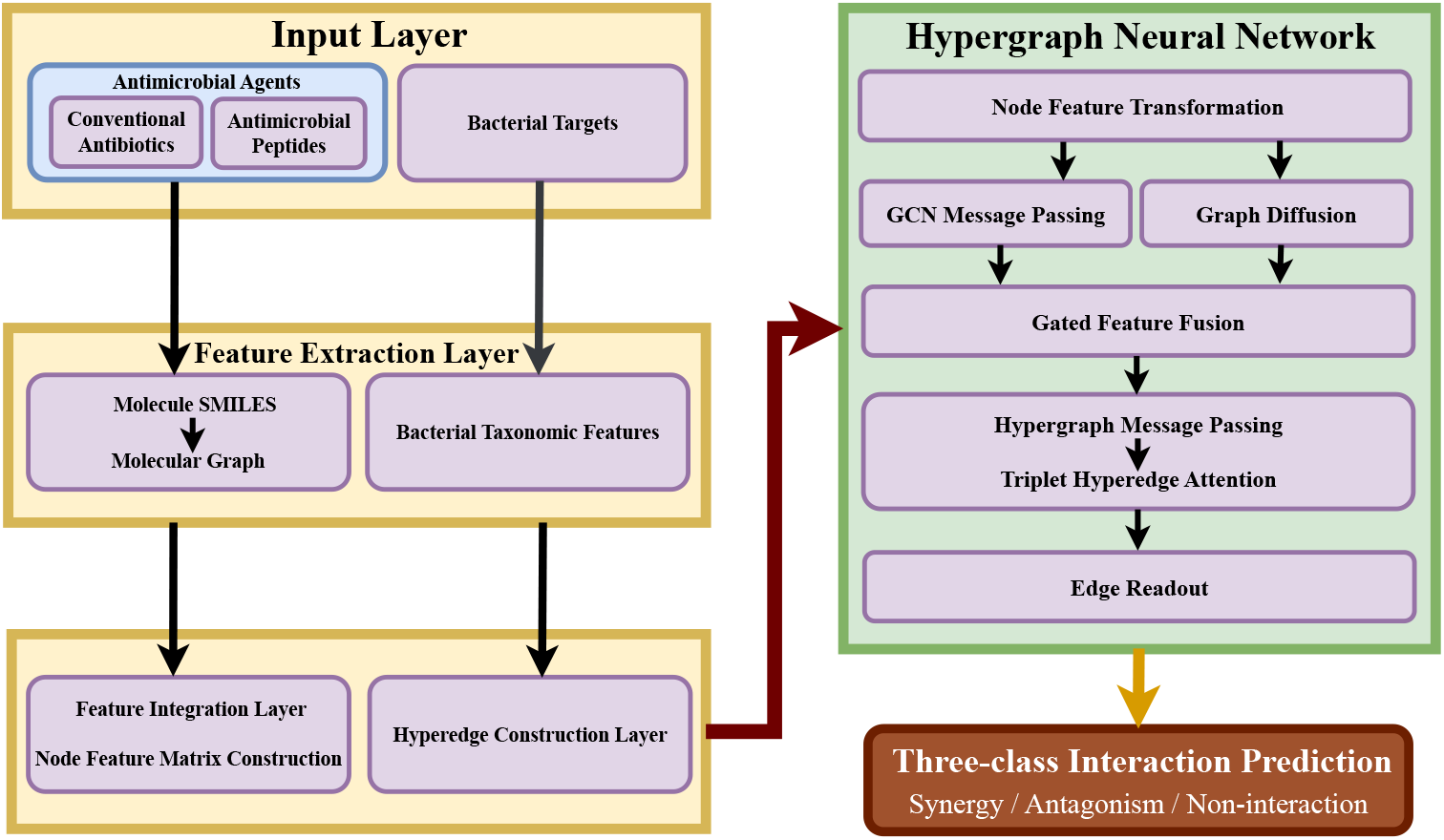
Overall workflow of the hypergraph-based antimicrobial interaction prediction framework. SMILES-derived molecular graph embeddings for antimicrobial agents and taxonomy-derived bacterial features are used to construct drug-drug-bacterium hyperedges. The HGNN combines graph convolution, graph diffusion, and hypergraph message passing, followed by a triplet readout layer that predicts synergy, antagonism, or non-interaction.

### 2.1 Data Acquisition and Preprocessing

The dataset integrated antimicrobial combination records involving conventional antibiotics and antimicrobial peptides (AMPs) tested against bacterial targets. Verified conventional antibiotic interaction data were obtained primarily from the Antibiotic Combination Database (ACDB) [35], whereas AMP-related activity and synergy records, including peptide-peptide and peptide-conventional antibiotic combinations, were obtained from the Database of Antimicrobial Activity and Structure of Peptides (DBAASP) [36].

Records were curated into antimicrobial agent-antimicrobial agent-bacterium triplets with experimentally reported interaction outcomes. Bacterial names were standardized and aligned to taxonomy-derived representations, and antimicrobial agents were stratified as AMPs or conventional antibiotics for pair-type-specific analyses.

### 2.2 Feature Engineering

Comprehensive feature engineering was performed to represent the distinct characteristics of antimicrobial agents and bacterial targets.

#### 2.2.1 Antimicrobial Agents Feature Engineering

Antimicrobial agents were primarily represented using their SMILES strings [37]. Each SMILES string was converted into a molecular graph compatible with PyTorch Geometric using RDKit [38]. For each atom in the molecule, a 10-dimensional feature vector was extracted. These features include normalized atomic number, degree, formal charge, explicit valence, an aromaticity flag, a ring membership flag, normalized total hydrogen count, normalized chiral tag, normalized hybridization state, and normalized implicit valence. Molecular connectivity was represented by bidirectional bond edges, which defined the graph structure used by the GCN-based molecular encoder [39].

As shown in Figure 2, these molecular graphs were then processed before HGNN training by a GCN-based molecular encoder, in which atom-level features were propagated according to the molecular bond-edge topology. The encoder projected 10-dimensional atomic features into a 64-dimensional hidden representation and applied three GCN blocks with Layer Normalization, GELU activation, dropout (*p* = 0.05), and residual connections in layers 2-3, producing a 128-dimensional molecular embedding.

**Figure 2.**
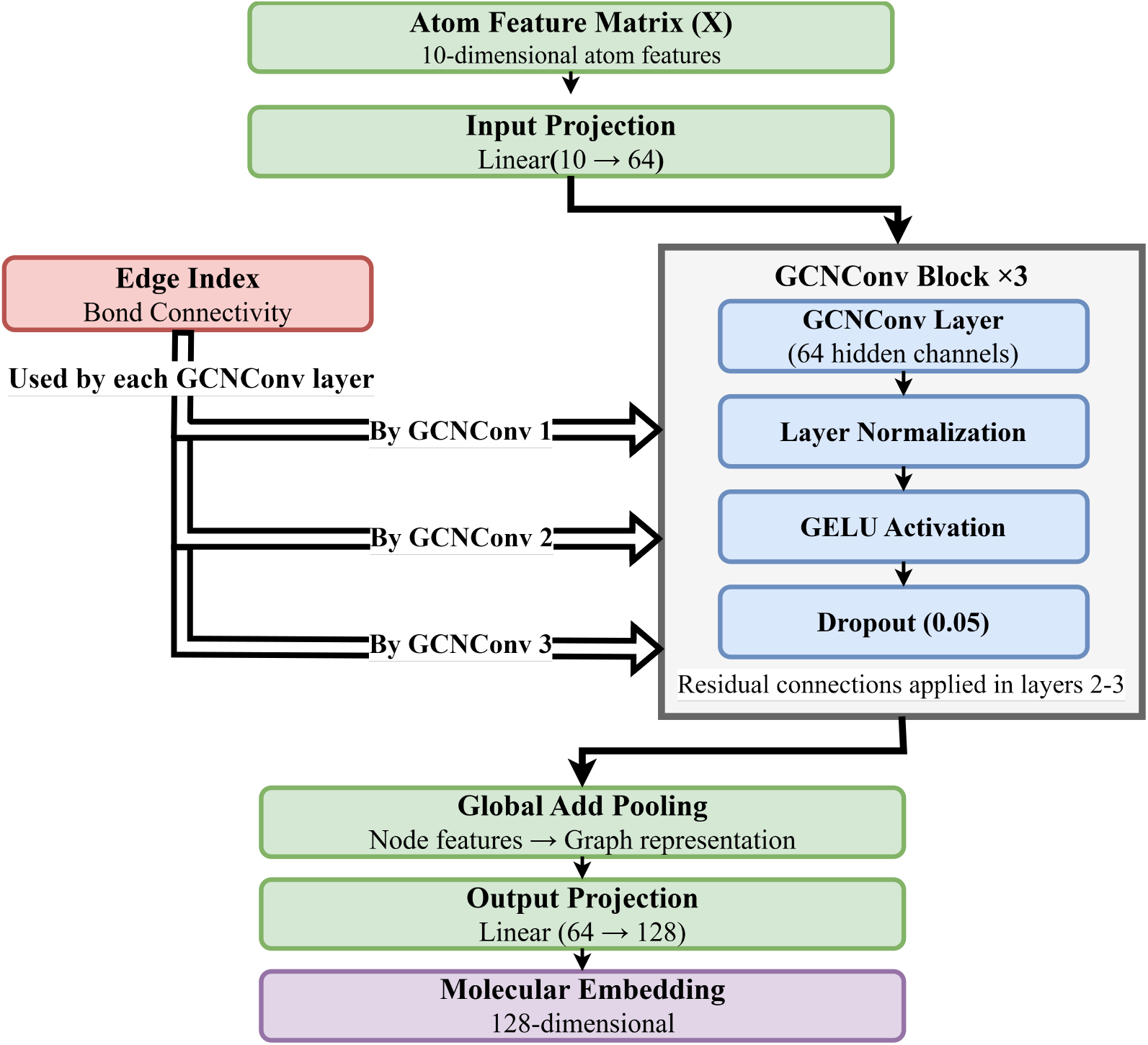
Molecular feature extraction using a GCN-based graph encoder. The workflow begins with atom-level features and edge indices, followed by a linear input projection. Three graph convolution layers sequentially apply graph convolution, Layer Normalization, GELU activation, dropout, and residual connections after the first layer [40]. Node representations are aggregated via Global Add Pooling to obtain a molecular-level graph embedding, which is linearly projected to yield the final molecular feature representation. Abbreviation: GELU, Gaussian Error Linear Unit.

#### 2.2.2 Bacterial Target Feature Engineering

To incorporate the evolutionary relationships among bacteria into our model, we developed a taxonomic distance embedding for each bacterium. This embedding captures the hierarchical structure of bacterial taxonomy, which includes genus, species, and strain levels [41], and quantifies the biological dissimilarity between different bacterial organisms into a matrix of shape (326, 326). For each of 326 bacterial nodes in the graph, its corresponding row, representing its taxonomic distances to all other bacteria in the dataset, was taken as its raw taxonomic feature vector. These taxonomic features were then normalized using Z-score normalization, by subtracting the mean and dividing by the standard deviation. In addition, the aligned taxonomic distance matrix was converted into a sparse similarity matrix used to create bacteria-bacteria message-passing edges. Distances were converted to similarities using exp(*−d/T*) with *T* = 2.0, followed by thresholding at 0.15 and top-*k* pruning with *k* = 12, as shown in Figure 3.

**Figure 3.**
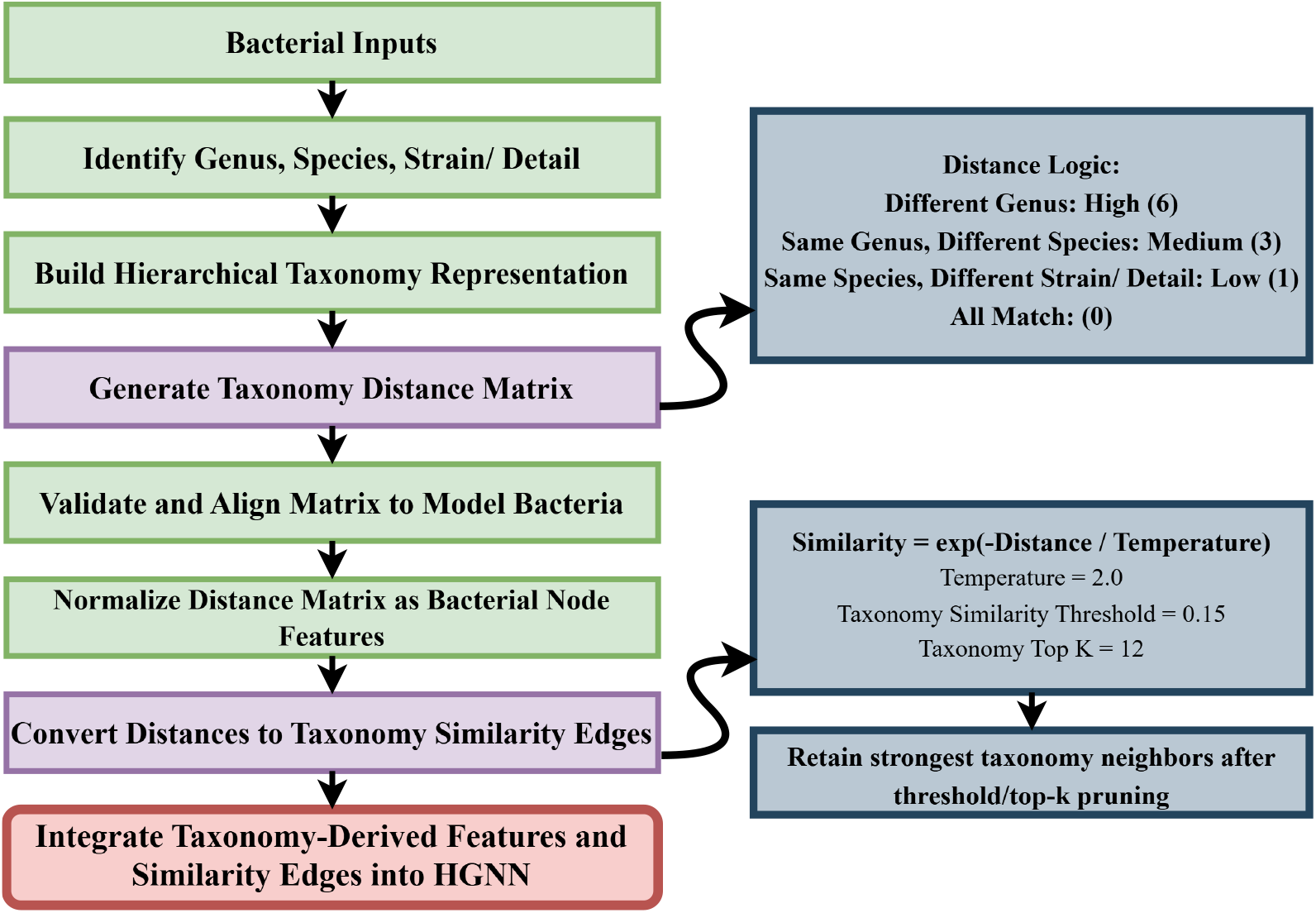
Construction of taxonomy-based bacterial feature representations. A hierarchical distance function is defined, with weights assigned to taxonomic levels to quantify phylogenetic proximity. Pairwise distances are computed to form a taxonomic distance matrix. The resulting matrix serves as a structured numerical representation of bacterial relationships for downstream modeling.

#### 2.2.3 Node Type Encoding and Final Feature Assembly

After molecular and bacterial features were generated, a two-dimensional node-type one-hot encoding was appended to distinguish antimicrobial-agent nodes from bacterial-target nodes. Drug nodes were represented by their 128-dimensional SMILES-derived molecular embedding, a drug-type indicator, and a zero-filled taxonomy vector. Bacterial nodes were represented by a zero-filled molecular embedding, a bacterial-type indicator, and the normalized taxonomy-derived feature vector. Concatenating these components produced the final 456-dimensional node feature matrix used by the HGNN.

### 2.3 Hypergraph Construction

Hyperedges were constructed to represent the three-way interactions. As shown in Figure 4, each hyperedge consists of three nodes: two corresponding to the antimicrobial agents and one corresponding to the bacterial target. The two antimicrobial nodes were sorted to form an unordered drug-pair representation, and the bacterial node was appended as the third element of the triplet. Observed interaction types of synergy, antagonism, and indifferent were mapped to numerical scores, and then mapped to the three model labels: synergy, antagonism, and non-interaction.

**Figure 4.**
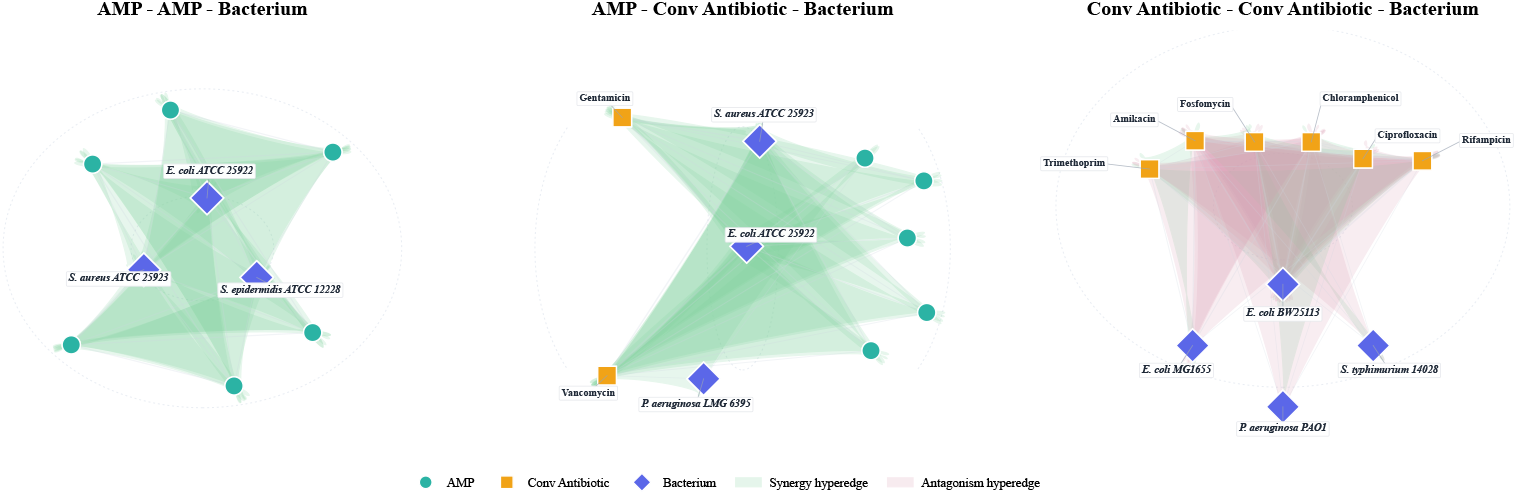
Visualization of sampled observed hyperedges across the three pair-type subsets. Each translucent shaded region represents one sampled three-node hyperedge consisting of two antimicrobial agents and one bacterial target. Node shapes and colors distinguish AMPs, conventional antibiotics, and bacterial targets, while shaded hyperedge colors denote the observed interaction class. Conv Antibiotic denotes conventional antibiotic.

### 2.4 Non-Interaction Class Construction

The non-interaction class was constructed from two sources: verified indifferent records in the dataset and synthetic non-interaction hyperedges generated by negative sampling. Verified indifferent samples were retained as observed non-interaction examples, whereas synthetic negatives represented unobserved Drug1-Drug2-Bacterium triplets generated only after the observed data had been split by unordered drug-pair groups. Synthetic negatives were added separately within each training, validation, and held-out test partition at a ratio of 25% relative to the observed triplets in that partition. Thus, synthetic negatives generated for validation or held-out testing were not reused as training samples. The negative sampling procedure used a mixed-curriculum strategy composed of 15% topology-aware negatives, 30% calibrated relation-preserving hard negatives, and 55% random negatives. Random negatives were generated by sampling two distinct antimicrobial nodes and one bacterial node while excluding all observed triplets. For topology-aware sampling, each candidate triplet was represented by the mean of its three node-feature vectors. The cosine distance between this candidate representation and sampled observed triplets was then calculated, and the candidate was retained only if its distance from every sampled observed triplet was at least 0.25 times the mean pairwise distance estimated among sampled observed triplets. Calibrated relation-preserving hard negatives followed the corrupted-triplet logic commonly used in relational representation learning: exactly one entity in an observed triplet was replaced by a similarity-ranked substitute of the same node type while preserving part of the observed relation structure [67, 68]. Drug replacements were sampled from a medium-similarity band after excluding the two closest drug neighbors, whereas bacterial replacements were sampled after excluding the closest bacterial neighbor. A candidate hard negative was retained only when it was absent from the observed graph, preserved exactly one observed sub-relation, and had no observed drug-bacterium sub-relation support.

### 2.5 Model Architecture

The core predictive model is a GCN-based hypergraph neural network designed to process these features and predict interaction outcomes.

#### 2.5.1 Initial Feature Transformation

The input node features of 456 dimensions first pass through a feature transformation module. This module consists of a linear layer projecting the input features to the model’s hidden channels, followed by Layer Normalization, a GELU activation function, and dropout [42–44].

#### 2.5.2 GCN Layers

The model employs a series of GCNConv layers for message passing [69]. The connectivity used by these layers is derived from a weighted adjacency matrix constructed from the triplet hyperedges using clique expansion, while allowing different contributions for drug-drug edges, drug-bacterium edges, and taxonomy-derived bacteria-bacteria edges. Self-loops are added and the matrix is symmetrically normalized [45]:

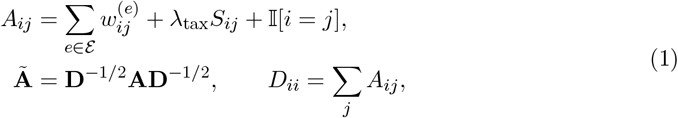

where 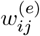 denotes the edge-type-specific contribution induced by hyperedge *e* and *S*_*ij*_ is the taxonomy-derived similarity between bacterial nodes. After the initial feature transformation, three graph convolution layers are applied, each followed by Layer Normalization and GELU activation, with residual connections added after the first layer.

#### 2.5.3 Graph Diffusion

As a complementary pathway, a Graph Diffusion module processes the transformed node features using the same normalized adjacency matrix. While the GCN message-passing branch learns local neighborhood representations, the diffusion branch propagates node information over the weighted graph topology to capture broader relational context among antimicrobial agents and bacterial targets. In this study, graph diffusion was performed for three propagation iterations. This propagation is implemented using an APPNP-like update inspired by personalized PageRank, in which each iteration computes a weighted combination of neighborhood-aggregated features and the initial transformed node representation, controlled by the propagation coefficient [70]:

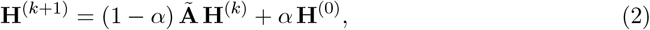

where *α* is initialized to 0.2 and learned during training.

#### 2.5.4 Hypergraph Message Passing

To preserve ternary context beyond pairwise expansion, a lightweight hypergraph block performs node-to-hyperedge-to-node propagation. For each triplet, a hyperedge representation is computed from the joint statistics of its three constituent nodes and then redistributed back to the incident nodes. The outputs of the graph convolution pathway, the diffusion pathway, and the hypergraph pathway are fused through learnable gates to form the final node representations, and the hyperedge context generated by the hypergraph block is projected and added to the triplet embedding before readout.

#### 2.5.5 Triplet Hyperedge Attention

For each three-node hyperedge, the message-passed representations of its constituent nodes are fed into a Triplet Hyperedge Attention module. It includes a triplet Multi-Layer Perceptron (MLP) that processes the concatenated features of all three nodes, node-specific attention to weigh individual node contributions, pairwise MLPs for drug-drug and drug-bacteria interactions, and an integration layer. This hierarchical approach decomposes the triplet interaction, separately modeling drug-drug and drug-bacteria sub-interactions before integrating them. Formally, given node embeddings **h**_1_, **h**_2_, **h**_3_ (Drug 1, Drug 2, Bacterium), the hyperedge embedding is computed as:

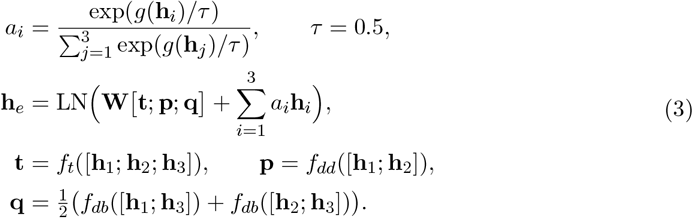

The output is a hyperedge embedding, further processed by Layer Normalization and dropout.

#### 2.5.6 Output Readout

The resulting hyperedge embedding is passed to an Edge Readout module for classification. This module has a shared MLP component followed by a 3-class head producing logits for synergy, antagonism, and non-interaction, an antagonism detector head producing a single auxiliary logit specifically for antagonism detection, and an interaction detector head producing a single auxiliary logit for discriminating real interactions from non-interactions. This multi-task learning framework enhances robustness by optimizing a shared representation for the main 3-class task together with targeted auxiliary objectives, acting as a regularizer to improve discrimination of the minority antagonism class and the non-interaction boundary [46].

### 2.6 Training Methodology

The main classification objective was a fold-specific weighted cross-entropy loss with class-specific boosts and label smoothing. Let **z**_*n*_ ∈ ℝ^3^ denote the 3-class logits for sample *n*, **p**_*n*_ = softmax(**z**_*n*_) the corresponding predicted class probabilities, and *y*_*n*_ ∈ {0, 1, 2} the class label (synergy, antagonism, non-interaction). With fold-specific class weights *α*_*c*_, class boosts *b*_*c*_, and label smoothing coefficient *ϵ* = 0.02, the smoothed target for class *c* is denoted by *q*_*n,c*_. The main classification loss is:

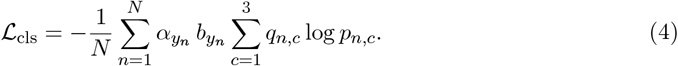

An auxiliary antagonism detector was trained, and additional auxiliary objectives were used to improve separation between interaction and non-interaction classes and to strengthen class margins. The total training loss is:

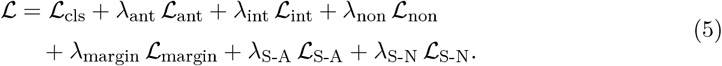

Here, ℒ_ant_, ℒ_int_, and ℒ_non_ denote auxiliary binary cross-entropy losses for antagonism detection, interaction detection, and non-interaction separation, respectively, while the remaining terms impose class-margin and synergy-specific pairwise penalties.

Initial class weights were computed separately for each training fold and adjusted. Verified indifferent samples remained active from the start of training as observed non-interaction examples, whereas synthetic negatives were introduced gradually using a curriculum schedule after a short warm-up. Mild epoch-level upsampling was applied only to the antagonism class. Optimization uses AdamW (initial learning rate=0.0002, weight decay=0.00001, AMSGrad) with a ReduceLROnPlateau scheduler (factor=0.5, patience=4, threshold=0.002, minimum learning rate =5% of the initial rate) [47–49]. The 4,391 observed samples were split by unordered drug pair into train/validation (3,521 edges) and held-out test (870 edges). A 5-fold grouped cross-validation was then performed on the train/validation portion, with additional synthetic non-interaction samples generated at 25% of the observed fold size for both training and validation [50]. Mini-batches were stratified (size=64, *≥* 4 samples/class where possible), gradients were clipped (norm=1.0) [51], and training ran for up to 100 epochs with early stopping (patience=10) [52]. Exponential moving average weights (decay=0.995) were used for validation and checkpoint selection. To avoid structural leakage, adjacency matrices for training, validation, and held-out testing were constructed only from positive training edges.

### 2.7 Evaluation Metrics and Validation Strategy

Model performance was assessed using accuracy, weighted F1 score, macro F1 score, precision, recall, ROC-AUC [53], and a model-specific weighted error index. Per-class accuracies for synergy, antagonism, and non-interaction, as well as balanced accuracy, were also computed. The weighted error index (WEI) was defined as a weighted class-level error rate::

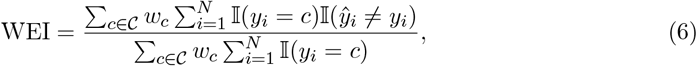

where *C* = {synergy, antagonism, non-interaction}, *y*_*i*_ and ŷ _*i*_ denote the true and predicted labels of sample *i*, respectively, and *w*_*c*_ denotes the class-specific penalty weight. In this study, the weights were set to *w*_synergy_ = 2.0, *w*_antagonism_ = 1.5, and *w*_non-interaction_ = 0.5, assigning greater penalty to errors in the experimentally observed interaction classes, particularly synergy. Lower CI values therefore indicate better weighted class-level prediction performance. Both validation and final held-out testing were performed in the same three-class setting, where the non-interaction class comprised verified indifferent samples together with synthetic negatives generated at the predefined ratio. Model selection during cross-validation was based on a composite score emphasizing balanced three-class performance while preserving the two real interaction classes.

### 2.8 Prospective Synergy Candidate Generation

After model training and held-out evaluation, the selected GCN-based HGNN checkpoint was used for prospective antimicrobial-combination generation. For each target bacterium, candidate antimicrobial pairs were assembled from the antimicrobial-agent node set using a bacterium-specific candidate-pool strategy. This strategy prioritized agents with direct observed support against the target bacterium, agents supported by taxonomically related bacteria, and globally frequent antimicrobial agents in the observed training interaction graph. Candidate unordered antimicrobial pairs were then combined with the target bacterium to form query hyperedges of the form Drug1-Drug2-Bacterium.

During inference, each query hyperedge was scored by the trained model using the observed training interaction graph to construct the prediction adjacency matrix. The query candidate batch itself was not used to define the support topology, preventing artificial coupling among candidate predictions. For each candidate pair, the model produced a three-class probability distribution over synergy, antagonism, and non-interaction. The primary generation metric was the predicted synergy probability,

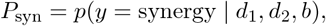

where *d*_1_ and *d*_2_ denote the two antimicrobial agents and *b* denotes the bacterial target. Prediction uncertainty was estimated from the normalized entropy of the three-class softmax distribution:

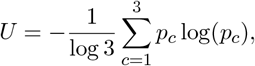

where *U ∈* [0, 1], with lower values indicating more confident predictions. Candidates were initially ranked using an uncertainty-adjusted score,

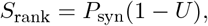

followed by diversity-aware re-ranking to avoid over-representation of a single antimicrobial agent among the top predictions.

To distinguish rediscovered interactions from genuinely prospective predictions, a strict source-novelty audit was performed against the complete training dataset derived from ACDB and DBAASP [35, 36]. A candidate was considered source-novel only if the exact antimicrobial pair was absent from the training data across all bacterial species. Final candidates were selected by combining source-novelty, *P*_syn_, uncertainty, and antimicrobial-pair class.

### 2.9 Ablation Study Construction

To evaluate the contribution of the GCN-based architecture, we constructed a set of ablation and baseline models under the same three-class prediction framework. The proposed model was the GCN-based HGNN implementation. Neural ablations replaced the main graph message-passing backbone with alternative operators, including TransformerConv, GATConv, and GINE-based message passing [71–74], while preserving the same curated dataset, node feature construction, drug-pair-grouped train/test split, negative sampling ratio, three-class labels, and held-out evaluation protocol. In addition, two classical machine learning baselines were evaluated: Random Forest and multinomial logistic regression [82, 83]. These baselines replaced the neural graph backbone with fixed triplet-level feature representations derived from the same node attributes and support-graph topology, including antimicrobial-pair summaries, antimicrobial-bacterium relation features, and graph-derived support statistics. All ablation and baseline models were evaluated on the held-out three-class test setting using the same primary metrics as the GCN-based model.

## 3 Results

This section reports the performance of the GCN-based HGNN in three-class cross-validation and drug-pair-grouped held-out evaluation.

### 3.1 Dataset and Preprocessing Outcomes Summary

After curation and taxonomy alignment, the final observed dataset contained 4,391 verified antimicrobial agent-antimicrobial agent-bacterium triplets labeled as synergy, antagonism, or indifferent. These triplets involved 571 antimicrobial agents, including 121 AMPs, and 326 bacterial targets. The observed label distribution was 3,484 synergy, 739 antagonism, and 168 verified indifferent interactions.

For pair-type-specific analyses, the dataset was stratified into 415 AMP-AMP-bacterium, 326 AMP-conventional antibiotic-bacterium, and 3,650 conventional antibiotic-conventional antibiotic-bacterium triplets. The dataset included 2,251 unique observed unordered drug-pair groups: 115 AMP-AMP, 180 AMP-conventional antibiotic, and 1,956 conventional antibiotic-conventional antibiotic pairs. In the observed dataset, AMP-containing triplets were limited to synergy labels, whereas antagonistic and verified indifferent records occurred only in the conventional antibiotic-conventional antibiotic subset.

The preprocessing pipeline generated 128-dimensional SMILES-derived molecular embeddings for antimicrobial agents, taxonomy-derived bacterial representations, and a final 456-dimensional node feature matrix for HGNN input. Using unordered drug-pair-grouped splitting, the observed triplets were divided into 3,521 train/validation triplets and 870 held-out triplets. To support three-class evaluation, 217 synthetic presumed non-interaction triplets were added to the held-out partition, resulting in a final held-out evaluation set of 1,087 instances.

### 3.2 Performance of the GCN-Based HGNN Model

The GCN-based HGNN model was evaluated across 5 grouped cross-validation folds. Table 1 summarizes the metrics of the best checkpoint selected within each fold. Fold 3 showed the best overall validation performance and was selected for held-out evaluation, with a selection score of 0.65.

**Table 1.**
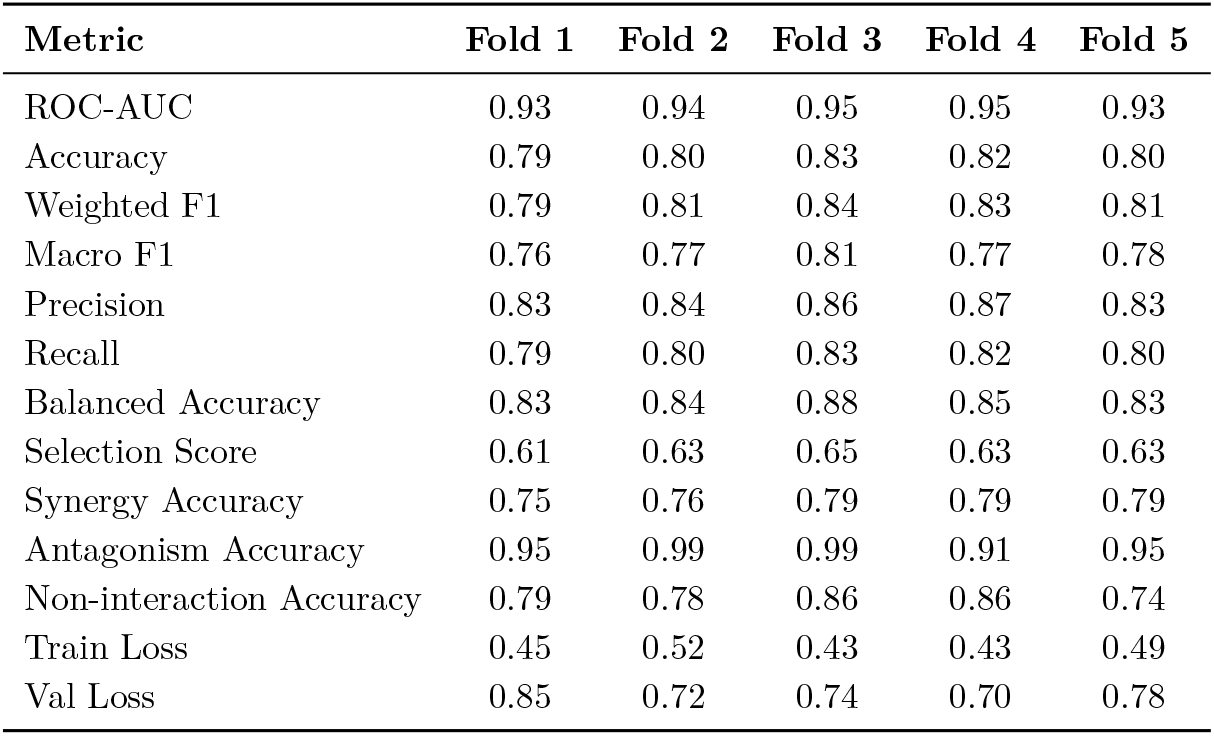
Cross-validation performance of the selected validation model state in each fold for the GCN-based HGNN model.

| Metric | Fold 1 | Fold 2 | Fold 3 | Fold 4 | Fold 5 |
| --- | --- | --- | --- | --- | --- |
| ROC-AUC | 0.93 | 0.94 | 0.95 | 0.95 | 0.93 |
| Accuracy | 0.79 | 0.80 | 0.83 | 0.82 | 0.80 |
| Weighted F1 | 0.79 | 0.81 | 0.84 | 0.83 | 0.81 |
| Macro F1 | 0.76 | 0.77 | 0.81 | 0.77 | 0.78 |
| Precision | 0.83 | 0.84 | 0.86 | 0.87 | 0.83 |
| Recall | 0.79 | 0.80 | 0.83 | 0.82 | 0.80 |
| Balanced Accuracy | 0.83 | 0.84 | 0.88 | 0.85 | 0.83 |
| Selection Score | 0.61 | 0.63 | 0.65 | 0.63 | 0.63 |
| Synergy Accuracy | 0.75 | 0.76 | 0.79 | 0.79 | 0.79 |
| Antagonism Accuracy | 0.95 | 0.99 | 0.99 | 0.91 | 0.95 |
| Non-interaction Accuracy | 0.79 | 0.78 | 0.86 | 0.86 | 0.74 |
| Train Loss | 0.45 | 0.52 | 0.43 | 0.43 | 0.49 |
| Val Loss | 0.85 | 0.72 | 0.74 | 0.70 | 0.78 |

### 3.3 Held-Out Test Set Results of the GCN-Based HGNN Model

The checkpoint selected from Fold 3 was evaluated on the held-out three-class test set. As shown in Table 2, the model achieved an accuracy of 0.83, weighted F1-score of 0.84, macro F1-score of 0.80, precision of 0.86, recall of 0.83, ROC-AUC of 0.95, and test loss of 0.48. Per-class performance showed that antagonism was identified most accurately (0.92), followed by non-interaction (0.85) and synergy (0.80).

**Table 2.** Held-out three-class test performance of the selected GCN-based HGNN model.

| Metric | Value |
| --- | --- |
| Test Loss | 0.48 |
| Accuracy | 0.83 |
| Weighted F1 | 0.84 |
| Macro F1 | 0.80 |
| Precision | 0.86 |
| Recall | 0.83 |
| ROC-AUC | 0.95 |
| WEI | 0.18 |
| Synergy Accuracy | 0.80 |
| Antagonism Accuracy | 0.92 |
| Non-interaction Accuracy | 0.85 |

Performance varied across interaction pair types (Table 3). Among AMP-containing combinations, the strongest overall performance was observed for AMP-AMP pairs, with an accuracy of 0.89 and weighted F1-score of 0.89. AMP-conventional antibiotic pairs also showed strong performance, with an accuracy of 0.85 and macro F1-score of 0.83. Conventional antibiotic-conventional antibiotic pairs formed the largest and most label-diverse subset and were the only pair type containing held-out antagonism examples; performance in this subset remained robust, with an accuracy of 0.82, weighted F1-score of 0.83, and macro F1-score of 0.79.

**Table 3.**
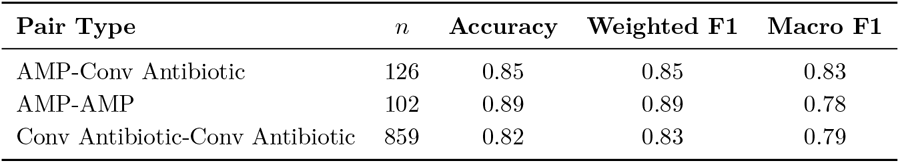
Held-out test performance of the selected GCN-based HGNN model stratified by interaction pair type. In this table, Conv Antibiotic denotes conventional antibiotic.

| Pair Type | $n$ | Accuracy | Weighted F1 | Macro F1 |
| --- | --- | --- | --- | --- |
| AMP-Conv Antibiotic | 126 | 0.85 | 0.85 | 0.83 |
| AMP-AMP | 102 | 0.89 | 0.89 | 0.78 |
| Conv Antibiotic-Conv Antibiotic | 859 | 0.82 | 0.83 | 0.79 |

### 3.4 Prospective Source-Novel Synergy Candidates for *Pseudomonas aeruginosa*

After held-out evaluation, the trained GCN-based HGNN was used to generate prospective synergy predictions for *Pseudomonas aeruginosa*. Candidate pairs were filtered using a strict source-novelty audit against the ACDB/DBAASP-derived training dataset [35,36]. Accordingly, the interactions reported in Table 4 were not present as exact antimicrobial pairs in the training data used in this study. This filtering step was applied to reduce direct rediscovery of source-data interactions.

**Table 4.**
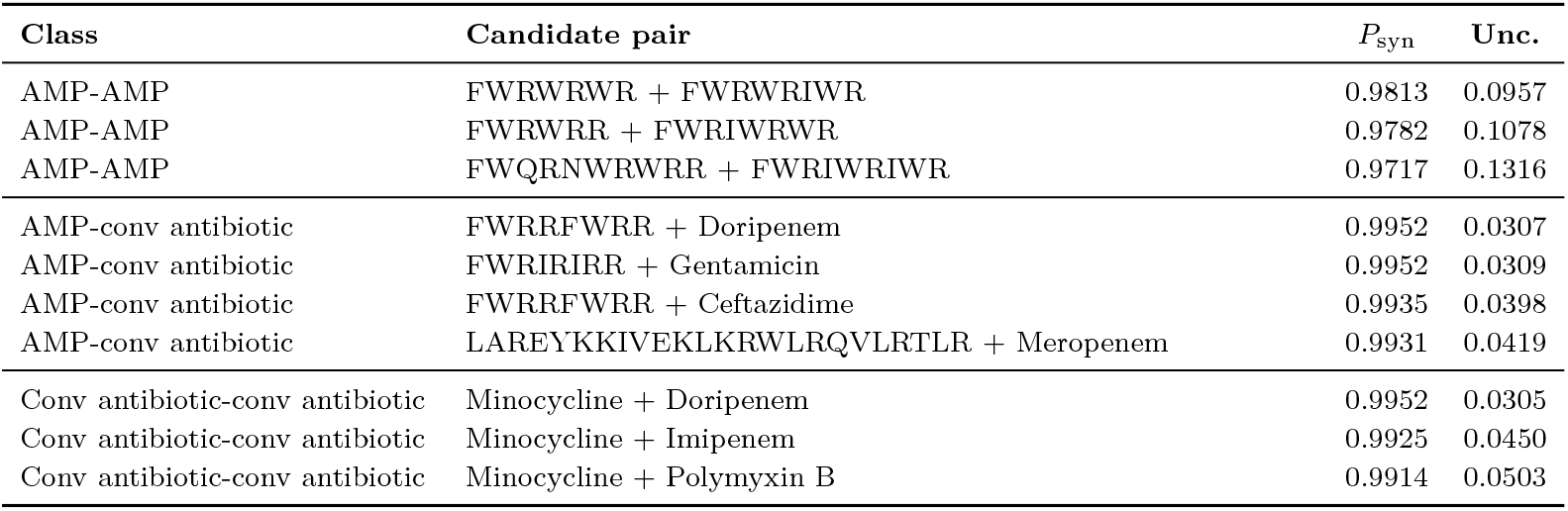
Top ten novel predicted synergy candidates for *Pseudomonas aeruginosa*. All listed interactions were absent from the ACDB/DBAASP-derived training data for Pseudomonas aeruginosa. Candidates were selected based on predicted synergy probability and prediction uncertainty. “Conv antibiotic” denotes conventional antibiotic.

The final top-ten list included representatives from all three antimicrobial-pair classes: three AMP-AMP pairs, four AMP-conventional antibiotic pairs, and three conventional antibiotic-conventional antibiotic pairs.

### 3.5 Ablation Study

To assess the contribution of the proposed GCN-based HGNN architecture, we compared it with five alternative predictive architectures implemented under the same three-class training and held-out evaluation framework: Transformer, GAT, GINE, Random Forest, and multinomial logistic regression. All alternative models used the same hypergraph-derived triplet representation, allowing the comparison to focus on the effect of the predictive architecture rather than differences in input formulation. As shown in Figure 5, the GCN-based model achieved the strongest overall held-out performance across all primary metrics (Table 5). Among the alternative architectures, the closest models were the Transformer and GINE variants, both of which reached a weighted F1-score of 0.80, whereas Random Forest and multinomial logistic regression showed lower overall performance.

**Table 5.**
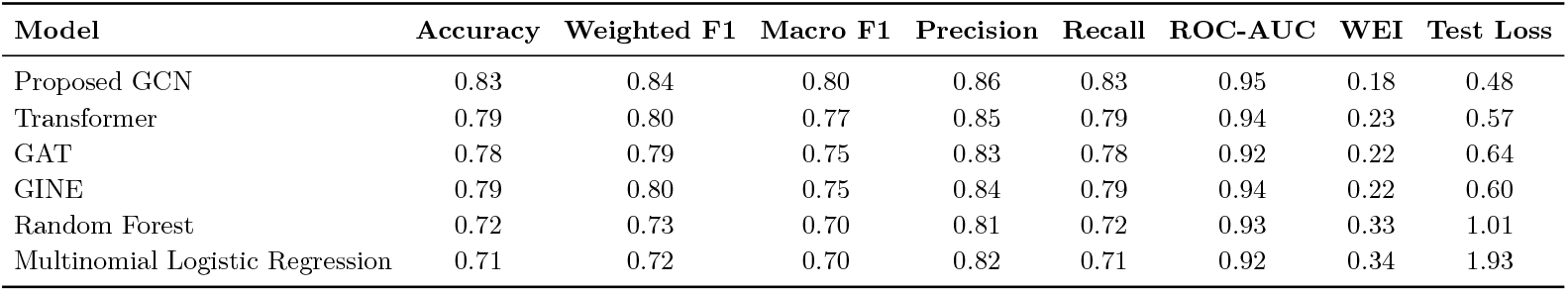
Held-out three-class test performance of the GCN-based HGNN model and alternative predictive architectures.

| Model | Accuracy | Weighted F1 | Macro F1 | Precision | Recall | ROC-AUC | WEI | Test Loss |
| --- | --- | --- | --- | --- | --- | --- | --- | --- |
| Proposed GCN | 0.83 | 0.84 | 0.80 | 0.86 | 0.83 | 0.95 | 0.18 | 0.48 |
| Transformer | 0.79 | 0.80 | 0.77 | 0.85 | 0.79 | 0.94 | 0.23 | 0.57 |
| GAT | 0.78 | 0.79 | 0.75 | 0.83 | 0.78 | 0.92 | 0.22 | 0.64 |
| GINE | 0.79 | 0.80 | 0.75 | 0.84 | 0.79 | 0.94 | 0.22 | 0.60 |
| Random Forest | 0.72 | 0.73 | 0.70 | 0.81 | 0.72 | 0.93 | 0.33 | 1.01 |
| Multinomial Logistic Regression | 0.71 | 0.72 | 0.70 | 0.82 | 0.71 | 0.92 | 0.34 | 1.93 |

**Figure 5.**
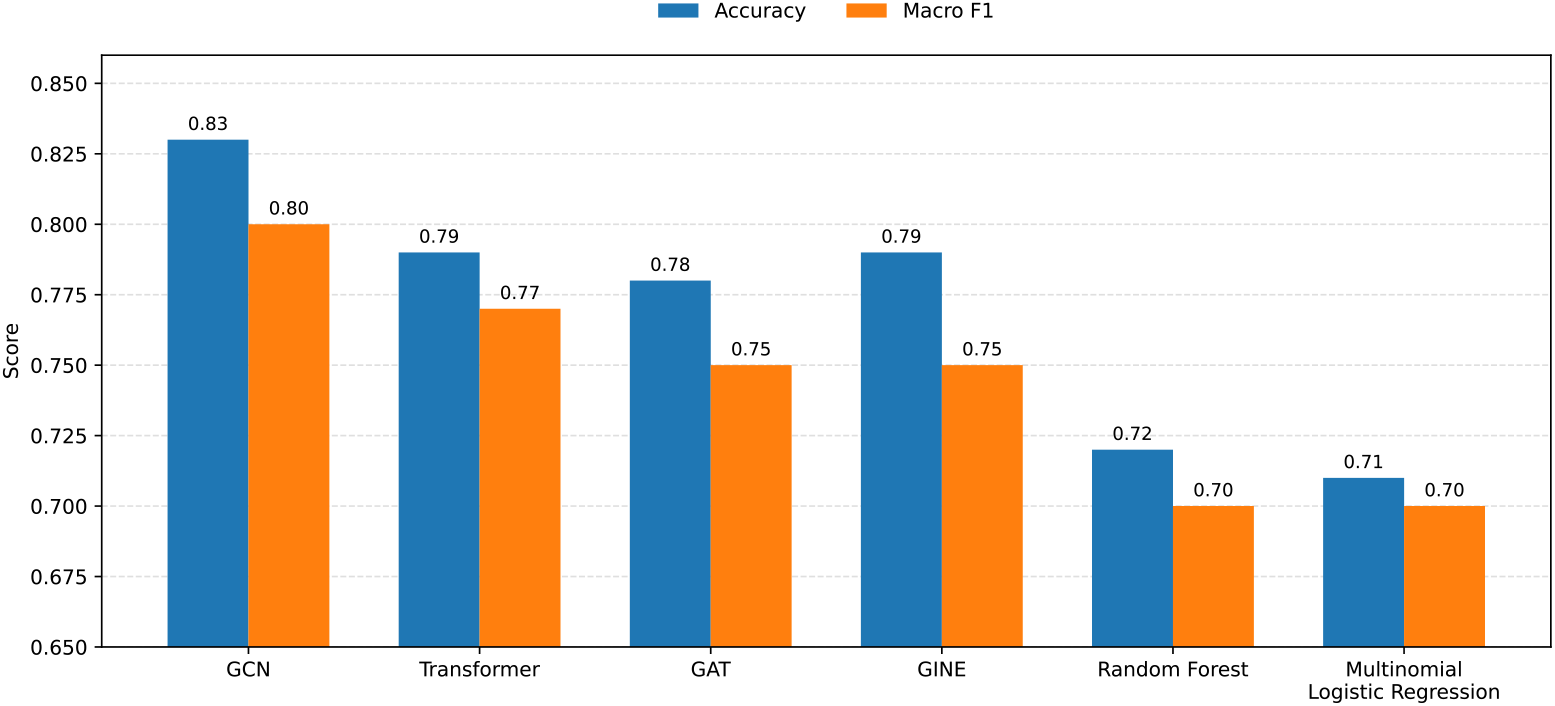
Comparison of held-out test accuracy and macro F1 across the original GCN-based HGNN model, neural backbone ablations, and classical baselines.

## 4 Discussion

A key advantage of the HGNN framework lies in its ability to encode expanding antimicrobial combination datasets using explicit ternary hyperedges. The hypergraph structure directly represents each observed Drug1-Drug2-Bacterium interaction as a compact hyperedge [45]. This design supports batched evaluation of large numbers of candidate combinations and is therefore well suited for prioritizing AMP-containing combinations and conventional antibiotic-conventional antibiotic combinations across diverse bacterial panels.

The model’s performance supports the value of integrating molecular, taxonomic, pairwise-graph, and hypergraph-level information. The GCN and diffusion branches capture local and broader graph structure, whereas the hypergraph pathway preserves the ternary Drug1-Drug2-Bacterium context that is central to antimicrobial combination effects. This design helps maintain performance across heterogeneous antimicrobial agents and bacterial targets, with its contribution further supported by the ablation results.

Recent antimicrobial-combination prediction studies differ in scope, labels, input features, and validation protocols, limiting strict head-to-head comparison. As summarized in Table 6, our proposed GCN-HGNN reported higher accuracy and ROC-AUC than the two contextual benchmarks while extending the task to ternary antimicrobial agent-antimicrobial agent-bacterium hyperedges and three-class prediction.

**Table 6.**
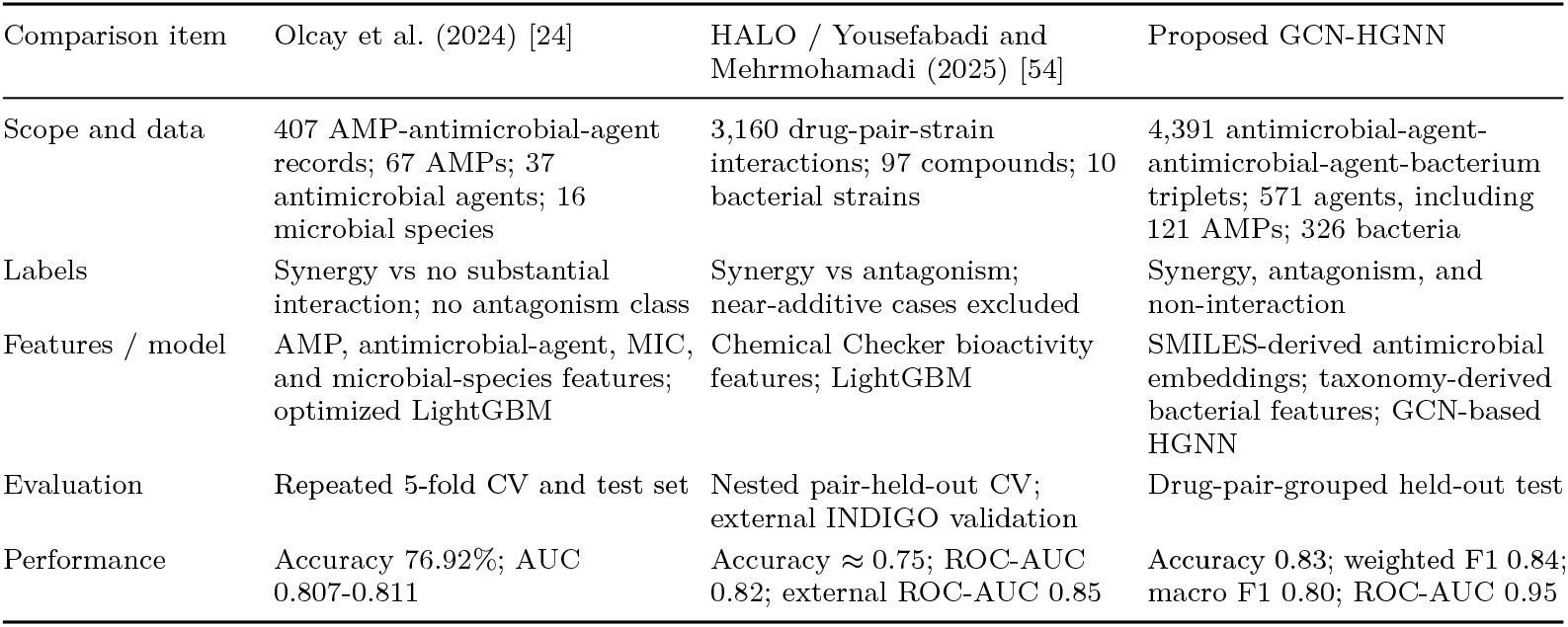
Comparison with antimicrobial-combination prediction studies.

| Comparison item | Olcay et al. (2024) [24] | HALO / Yousefabadi and Mehrmohamadi (2025) [54] | Proposed GCN-HGNN |
| --- | --- | --- | --- |
| Scope and data | 407 AMP-antimicrobial-agent records; 67 AMPs; 37 antimicrobial agents; 16 microbial species | 3,160 drug-pair-strain interactions; 97 compounds; 10 bacterial strains | 4,391 antimicrobial-agent-antimicrobial-agent-bacterium triplets; 571 agents, including 121 AMPs; 326 bacteria |
| Labels | Synergy vs no substantial interaction; no antagonism class | Synergy vs antagonism; near-additive cases excluded | Synergy, antagonism, and non-interaction |
| Features / model | AMP, antimicrobial-agent, MIC, and microbial-species features; optimized LightGBM | Chemical Checker bioactivity features; LightGBM | SMILES-derived antimicrobial embeddings; taxonomy-derived bacterial features; GCN-based HGNN |
| Evaluation | Repeated 5-fold CV and test set | Nested pair-held-out CV; external INDIGO validation | Drug-pair-grouped held-out test |
| Performance | Accuracy 76.92%; AUC 0.807-0.811 | Accuracy ≈ 0.75; ROC-AUC 0.82; external ROC-AUC 0.85 | Accuracy 0.83; weighted F1 0.84; macro F1 0.80; ROC-AUC 0.95 |

Pair-type stratification offers a more informative view of model behavior than aggregate performance alone. In the evaluated held-out set, the AMP-containing subsets, including AMP-AMP and AMP-conventional antibiotic combinations, were much smaller than the conventional antibiotic-conventional antibiotic subset, whereas conventional antibiotic-Conventional antibiotic combinations dominated the test distribution and were the only subset containing held-out antagonistic cases in addition to non-interaction examples. This makes the conventional antibiotic-conventional antibiotic setting both larger and label-wise more demanding, while results for AMP-AMP pairs should be interpreted cautiously because that subset was smaller and lacked held-out antagonistic cases. This pattern was also visible in the learned triplet embedding space (Figure 6). In the pair-type-stratified projection, AMP-AMP and AMP-conventional antibiotic triplets occupied smaller regions of the learned space and contained only synergistic records, whereas conventional antibiotic-conventional antibiotic triplets covered a broader region and included synergy, antagonism, and indifferent labels. Antagonistic records formed a more localized region within the conventional antibiotic-conventional antibiotic panel, while indifferent records partially overlapped with both synergistic and antagonistic regions. Therefore, the projection supports the interpretation that pair-type-specific patterns are influenced by label availability, subset size, and the structure of the learned representation.

**Figure 6.**
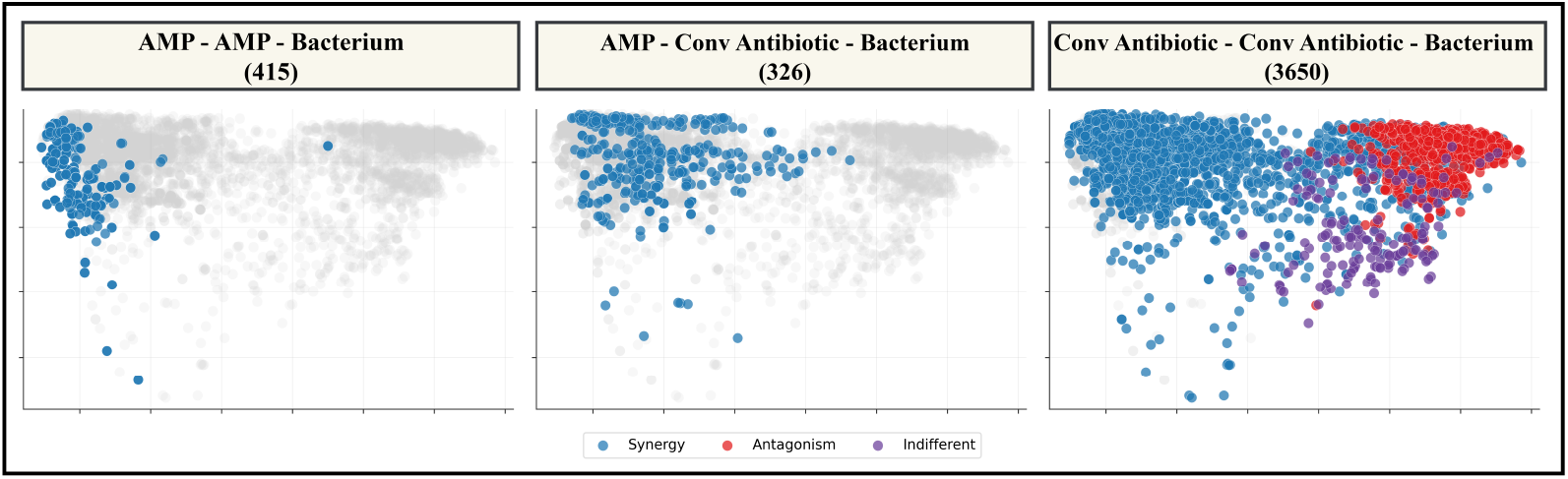
Pair-type-stratified PCA [55] projection of learned triplet embeddings from the selected GCN-based HGNN model. Each panel highlights one interaction pair type, while all triplets are shown in gray as background. Points are colored by interaction label. AMP-AMP and AMP-conventional antibiotic triplets contain only synergistic records in the curated dataset, whereas conventional antibiotic-conventional antibiotic triplets include synergy, antagonism, and indifferent records.

A possible explanation is that, although both AMPs and conventional antibiotics were encoded through the same structure-based molecular feature pipeline, AMP-AMP combinations may be easier for the model to separate in this evaluation setting, whereas conventional antibiotic-conventional antibiotic and AMP-conventional antibiotic pairs involve broader interaction patterns that are harder to distinguish. Overall, these findings suggest that pair-type performance depends jointly on subset size, label composition, and the complexity of the underlying representation.

Among the source-novel *P. aeruginosa* candidates in Table 4, seven of the prioritized pairs involved AMPs, comprising three AMP-AMP and four AMP-antibiotic combinations. Most of the short peptide sequences in these pairs are cationic and W/R-rich, a physicochemical class expected to interact with anionic bacterial envelopes through electrostatic binding, interfacial partitioning, and membrane perturbation [56]. This makes the enrichment of sequences such as FWRWRWR, FWRWRIWR, FWRWRR, and related peptides biologically plausible rather than purely statistical. Prior high-throughput AMP studies show that small sequence substitutions and scrambling can markedly affect antipseudomonal activity [57, 58], while combinatorial studies indicate that synergy patterns among short cationic AMPs can be sequence- and partner-dependent in multidrug-resistant *P. aeruginosa* [76]. The AMP-AMP predictions should therefore be regarded as prospective but mechanistically credible: AMP combinations can be synergistic, and in some systems synergy depends on concentration ratio and aggregation state, with peptide co-aggregation influencing membrane disruption [21, 59]. The AMP-conventional antibiotic predictions have stronger external support. The W/R-rich peptide sequences `FWRRFWRR` and `FWRIRIRR` correspond to human-lactoferricin-derived LF11 peptides previously reported to display antipseudomonal activity against *P. aeruginosa* planktonic cultures and biofilms [75]. Their prioritization with doripenem, gentamicin, and ceftazidime is biologically plausible in light of broader evidence that short cationic AMPs and lactoferricin-derived peptides can synergize with conventional antibiotics against multidrug-resistant *P. aeruginosa* models *in vitro* and *in vivo*; however, the exact LF11-derived peptide-antibiotic pairs reported here remain prospective predictions requiring direct experimental validation [20,60,76]. Similarly, the peptide sequence LAREYKKIVEKLKRWLRQVLRTLR corresponds to P10, an LL-37-derived AMP for which synergistic or additive activity with ceftazidime or doripenem has been reported against resistant *P. aeruginosa* isolates; its predicted pairing with meropenem therefore represents a related carbapenem-containing candidate, although the exact P10-meropenem pair still requires direct validation [77]. In parallel, the conventional antibiotic-conventional antibiotic subset showed a second coherent pattern: all three pairs contained minocycline, combined with doripenem, imipenem, or polymyxin B. This is consistent with the resistance architecture of *P. aeruginosa*, where low outer-membrane permeability and multidrug efflux limit intracellular antibiotic accumulation [61–63, 81]. Among these three minocycline-containing predictions, minocycline-polymyxin B has the strongest pair-level support: outer-membrane perturbation can potentiate otherwise weakly active antibiotics in Gram-negative bacteria, permeability-enhancing adjuvants can potentiate minocycline or other tetracyclines against *P. aeruginosa*, and Olsson et al. reported enhanced or synergistic activity for polymyxin B-minocycline against multidrug-resistant *P. aeruginosa* [64, 65, *78, 80]*. Polymyxin B combinations with doripenem or meropenem have also shown enhanced activity against resistant *P. aeruginosa* [78, 79]. By contrast, minocycline-doripenem and minocycline-imipenem remain more prospective; available evidence supports only related *β*-lactam/tetracycline interaction contexts, not direct validation of these exact pairs [66]. Overall, Table 4 suggests that the model prioritized source-novel combinations around two coherent axes: AMP-mediated envelope disruption and antibiotic accumulation rescue.

## 5 Implications and Future Directions

The proposed framework can support experimental prioritization by ranking antimicrobial-agents and bacterium triplets for follow-up testing. Its main practical value is to reduce the number of combinations requiring empirical screening, particularly for AMP-containing combinations and conventional antibiotic pairs evaluated across multiple bacterial targets.

Building on this foundation, future enhancements could amplify the model’s impact through several key avenues: expanding datasets with additional antimicrobial agent combinations and resistant bacterial strains to improve generalization and accuracy; advancing AMP feature engineering by incorporating amino acid sequences, physicochemical properties, and predicted secondary/tertiary structures beyond SMILES representations; implementing interpretability techniques to clarify decision-making and key interaction drivers; conducting prospective experimental validation of predicted synergistic combinations via wet-lab assays; extending predictions to dynamic and temporal effects, such as time-kill dynamics or resistance emergence; integrating richer biological context, including drug targets, pathways, mechanisms of action, bacterial virulence factors, and host-pathogen interactions; and refining negative sampling strategies for more informative examples to enhance model robustness. These developments will strengthen the framework as a tool for supporting computational prioritization of antimicrobial combinations.

## 6 Conclusion

We propose a GCN-based HGNN framework that represents each Drug1-Drug2-Bacterium observation as a single hyperedge, enabling context-dependent prediction of combination outcomes across bacterial targets. The framework functions as an effective computational triage tool: it ranks candidate synergistic combinations, highlights those warranting experimental validation, and reduces the wet-lab search space. These results establish machine learning as a practical addition to the toolkit for accelerating antimicrobial combination discovery.

## Code and Data Availability

The datasets used in this study were derived from the public resources described in the Methods section. The processed datasets, source code, and model-related files are publicly available in the following GitHub repository: https://github.com/Farzad-Midjani/Hypergraph-Antimicrobial-combiantion.

## Abbreviations

AMP: Antimicrobial Peptide
AMR: Antimicrobial Resistance
HGNN: Hypergraph Neural Network
GNN: Graph Neural Network
ROC-AUC: Receiver Operating Characteristic - Area Under the Curve
PK/PD: Pharmacokinetic/Pharmacodynamic
QSAR: Quantitative Structure-Activity Relationship
DBAASP: Database of Antimicrobial Activity and Structure of Peptides
ACDB: Antibiotic Combination Database
SMILES: Simplified Molecular Input Line Entry System
MLP: Multi-Layer Perceptron
GELU: Gaussian Error Linear Unit
WEI: Weigted Error Index

## References

[1] Walsh TR, Gales AC, Laxminarayan R, Dodd PC (2023) Antimicrobial Resistance: Addressing a Global Threat to Humanity. PLoS Med 20(7): e1004264. doi:10.1371/journal.pmed.1004264.

[2] O’Neill J. Antimicrobial Resistance: Tackling a Crisis for the Health and Wealth of Nations: December 2014. Review on Antimicrobial Resistance; 2014.

[3] Antimicrobial Resistance Collaborators. Global burden of bacterial antimicrobial resistance in 2019: a systematic analysis. Lancet. 2022;399(10325):629–655. doi:10.1016/S0140-6736(21)02724-0.

[4] Salam MA, Al-Amin MY, Salam MT, Pawar JS, Akhter N, Rabaan AA, Alqumber MAA. Antimicrobial resistance: A growing serious threat for global public health. Healthcare (Basel). 2023;11(13):1946. doi:10.3390/healthcare11131946.

[5] Van Boeckel TP, Pires J, Silvester R, Zhao C, Song J, Criscuolo NG, Gilbert M, Bonhoeffer S, Laxminarayan R. Global trends in antimicrobial resistance in animals in low- and middle-income countries. Science. 2019;365(6459):eaaw1944. doi:10.1126/science.aaw1944.

[6] Tatem AJ, Rogers DJ, Hay SI. Global transport networks and infectious disease spread. Adv Parasitol. 2006;62:293–343. doi:10.1016/S0065-308X(05)62009-X.

[7] Hassing RJ, Alsma J, Arcilla MS, van Genderen PJ, Stricker BH, Verbon A. International travel and acquisition of multidrug-resistant Enterobacteriaceae: a systematic review. Euro Surveill. 2015;20(47):30074. doi:10.2807/1560-7917.ES.2015.20.47.30074.

[8] Frost I, Van Boeckel TP, Pires J, Craig J, Laxminarayan R. Global geographic trends in antimicrobial resistance: the role of international travel. J Travel Med. 2019;26(8):taz036. doi:10.1093/jtm/taz036.

[9] Hendriksen RS, Munk P, Njage P, van Bunnik B, McNally L, Lukjancenko O, et al. Global monitoring of antimicrobial resistance based on metagenomics analyses of urban sewage. Nat Commun. 2019;10(1):1124. doi:10.1038/s41467-019-08853-3.

[10] Lerminiaux NA, Cameron ADS. Horizontal transfer of antibiotic resistance genes in clinical environments. Can J Microbiol. 2019;65(1):34–44. doi:10.1139/cjm-2018-0275.

[11] Tang KWK, Millar BC, Moore JE. Antimicrobial resistance (AMR). Br J Biomed Sci. 2023;80:11387. doi:10.3389/bjbs.2023.11387.

[12] Blair JMA, Webber MA, Baylay AJ, Ogbolu DO, Piddock LJV. Molecular mechanisms of antibiotic resistance. Nat Rev Microbiol. 2015;13(1):42–51. doi:10.1038/nrmicro3380.

[13] Tao S, Chen H, Li N, Wang T, Liang W. The spread of antibiotic resistance genes in vivo model. Can J Infect Dis Med Microbiol. 2022;2022:3348695. doi:10.1155/2022/3348695.

[14] Cassini A, Högberg LD, Plachouras D, Quattrocchi A, Hoxha A, Simonsen GS, et al. Attributable deaths and disability-adjusted life-years caused by infections with antibiotic-resistant bacteria in the EU and the European Economic Area in 2015: a population-level modelling analysis. Lancet Infect Dis. 2019;19(1):56–66. doi:10.1016/S1473-3099(18)30605-4.

[15] Tallarida RJ. Quantitative methods for assessing drug synergism. Genes Cancer. 2011;2(11):1003–1008. doi:10.1177/1947601912440575.

[16] Bognár B, Spohn R, Lázár V. Drug combinations targeting antibiotic resistance. npj Antimicrob Resist. 2024;2(1):29. doi:10.1038/s44259-024-00047-2.

[17] Balouiri M, Sadiki M, Ibnsouda SK. Methods for in vitro evaluating antimicrobial activity: a review. J Pharm Anal. 2016;6(2):71–79. doi:10.1016/j.jpha.2015.11.005.

[18] Gaudereto JJ, Perdigão-Neto LV, Leite GC, Espinoza EPS, Martins RCR, Prado GVB, Rossi F, Guimarães T, Levin AS, Costa SF. Comparison of methods for the detection of in vitro synergy in multidrug-resistant gram-negative bacteria. BMC Microbiol. 2020;20(1):97. doi:10.1186/s12866-020-01756-0.

[19] Ma X, Wang Q, Ren K, Xu T, Zhang Z, Xu M, Rao Z, Zhang X. A review of antimicrobial peptides: structure, mechanism of action, and molecular optimization strategies. Fermentation. 2024;10(11):540. doi:10.3390/fermentation10110540.

[20] Taheri-Araghi S. Synergistic action of antimicrobial peptides and antibiotics: current understanding and future directions. Front Microbiol. 2024;15:1390765. doi:10.3389/fmicb.2024.1390765.

[21] Yu G, Baeder DY, Regoes RR, Rolff J. Combination effects of antimicrobial peptides. Antimicrob Agents Chemother. 2016;60(3):1717–1724. doi:10.1128/AAC.02434-15.

[22] Cantrell JM, Chung CH, Chandrasekaran S. Machine learning to design antimicrobial combination therapies: promises and pitfalls. Drug Discov Today. 2022;27(6):1639–1651. doi:10.1016/j.drudis.2022.04.006.

[23] O’Jeanson A, Nielsen EI, Friberg LE. A model-based evaluation of the pharmacokinetics-pharmacodynamics (PKPD) of avibactam in combination with ceftazidime. JAC-Antimicrobial Resistance. 2025;7(2):dlaf036. doi:10.1093/jacamr/dlaf036.

[24] Olcay B, Ozdemir GD, Ozdemir MA, Ercan UK, Guren O, Karaman O. Prediction of the synergistic effect of antimicrobial peptides and antimicrobial agents via supervised machine learning. BMC Biomed. Eng. 2024;6(1):1. doi:10.1186/s42490-024-00075-z.

[25] Pérez de la Lastra JM, Wardell SJT, Pal T, de la Fuente-Nunez C, Pletzer D. From Data to Decisions: Leveraging Artificial Intelligence and Machine Learning in Combating Antimicrobial Resistance-a Comprehensive Review. J Med Syst. 2024;48(1):71. doi:10.1007/s10916-024-02089-5.

[26] Reiser P, Neubert M, Eberhard A, Torresi L, Zhou C, Shao C, et al. Graph neural networks for materials science and chemistry. Commun. Mater. 2022;3(1):93. doi:10.1038/s43246-022-00315-6.

[27] Zhang Z, Chen L, Zhong F, Wang D, Jiang J, Zhang S, et al. Graph neural network approaches for drug-target interactions. Curr. Opin. Struct. Biol. 2022;73:102327. doi:10.1016/j.sbi.2021.102327.

[28] Brochado AR, Telzerow A, Bobonis J, Banzhaf M, Mateus A, Selkrig J, Bassler S, Zietek M, Savitski MM, Typas A. Species-specific activity of antibacterial drug combinations. Nature. 2018;559:259–263. doi:10.1038/s41586-018-0278-9.

[29] Tang P-C, Sánchez-Hevia DL, Westhoff S, Fatsis-Kavalopoulos N, Andersson DI. Within-species variability of antibiotic interactions in Gram-negative bacteria. mBio. 2024;15(3):e00196–24. doi:10.1128/mbio.00196-24.

[30] Cacace E, Kim V, Varik V, Knopp M, Tietgen M, Brauer-Nikonow A, et al. Systematic analysis of drug combinations against Gram-positive bacteria. Nat. Microbiol. 2023;8(11):2196–2212. doi:10.1038/s41564-023-01486-9.

[31] Gu Y, Zu J, Sun Y, Zhang L. HIG-Syn: a hypergraph and interaction-aware multigranularity network for predicting synergistic drug combinations. Bioinformatics. 2025;41(Suppl_1):i86–i95. doi:10.1093/bioinformatics/btaf215.

[32] Bai S, Zhang F, Torr PHS. Hypergraph convolution and hypergraph attention. Pattern Recognit. 2021;110:107637. doi:10.1016/j.patcog.2020.107637.

[33] Liu X, Song C, Liu S, Li M, Zhou X, Zhang W. Multi-way relation-enhanced hypergraph representation learning for anti-cancer drug synergy prediction. Bioinformatics. 2022;38(20):4782–4789. doi:10.1093/bioinformatics/btac579.

[34] Feng Y, You H, Zhang Z, Ji R, Gao Y. Hypergraph neural networks. In: Proceedings of the AAAI Conference on Artificial Intelligence (AAAI-19), Vol. 33, No. 01; 2019. p. 3558–3565. doi:10.1609/aaai.v33i01.33013558.

[35] Lv J, Liu G, Dong W, Ju Y, Sun Y. ACDB: an antibiotic combination database. Front. Pharmacol. 2022;13:869983. doi:10.3389/fphar.2022.869983.

[36] Pirtskhalava M, Amstrong AA, Grigolava M, Chubinidze M, Alimbarashvili E, Vishnepolsky B, et al. DBAASP v3: database of antimicrobial/cytotoxic activity and structure of peptides as a resource for development of new therapeutics. Nucleic Acids Res. 2021;49(D1):D288–D297. doi:10.1093/nar/gkaa991.

[37] Weininger D. SMILES, a chemical language and information system. 1. Introduction to methodology and encoding rules. J. Chem. Inf. Comput. Sci. 1988;28(1):31–36. doi:10.1021/ci00057a005.

[38] Landrum G, Tosco P, Kelley B, Rodriguez R, Cosgrove D, Vianello R sriniker, Gedeck P, Jones G, Kawashima E NadineSchneider, Nealschneider D tadhurst-cdd, Dalke A, Swain M, Cole B, Turk S, Savelev A, Maeder N, Vaucher A, Wójcikowski M, Faara H, Take I, Walker R, Scalfani VF, Probst D, Ujihara K, Pahl A, godin g, Lehtivarjo J. rdkit/rdkit: 2025_09_5 (Q3 2025) Release. doi:10.5281/zenodo.18428170.

[39] Fey M, Lenssen JE (2019) Fast graph representation learning with PyTorch Geometric. In: ICLR Workshop on Representation Learning on Graphs and Manifolds. arXiv:1903.02428. doi:10.48550/arXiv.1903.02428.

[40] He K, Zhang X, Ren S, Sun J. Deep residual learning for image recognition. In: Proceedings of the IEEE Conference on Computer Vision and Pattern Recognition (CVPR). 2016;770-778. doi:10.1109/CVPR.2016.90.

[41] Federhen S. The NCBI taxonomy database. Nucleic Acids Res. 2012;40(Database issue):D136–D143. doi:10.1093/nar/gkr1178.

[42] Ba JL, Kiros JR, Hinton GE. Layer normalization. arXiv:1607.06450. 2016. doi:10.48550/arXiv.1607.06450.

[43] Hendrycks D, Gimpel K. Gaussian error linear units (GELUs). arXiv:1606.08415. 2016. doi:10.48550/arXiv.1606.08415.

[44] Srivastava N, Hinton G, Krizhevsky A, Sutskever I, Salakhutdinov R. Dropout: a simple way to prevent neural networks from overfitting. J Mach Learn Res. 2014;15:1929–1958.

[45] Zhou D, Huang J, Schölkopf B. Learning with hypergraphs: clustering, classification, and embedding. In: Advances in Neural Information Processing Systems 19 (NeurIPS 2006). pp. 1601–1608.

[46] Caruana R. Multitask learning. Mach Learn. 1997;28:41–75. doi:10.1023/A:1007379606734.

[47] Loshchilov I, Hutter F. Decoupled weight decay regularization. In: International Conference on Learning Representations (ICLR 2019).

[48] Reddi SJ, Kale S, Kumar S. On the convergence of Adam and beyond. In: International Conference on Learning Representations (ICLR 2018).

[49] Kingma DP, Ba J. Adam: a method for stochastic optimization. In: International Conference on Learning Representations (ICLR 2015).

[50] Kohavi R. A study of cross-validation and bootstrap for accuracy estimation and model selection. In: Proceedings of the Fourteenth International Joint Conference on Artificial Intelligence (IJCAI’95), Vol 2. 1995. p. 1137—1145.

[51] Pascanu R, Mikolov T, Bengio Y. On the difficulty of training recurrent neural networks. In: Proceedings of the 30th International Conference on Machine Learning. Proc Mach Learn Res. 2013;28(3):1310–1318.

[52] Prechelt L. Automatic early stopping using cross validation: quantifying the criteria. Neural Netw. 1998;11(4):761–767. doi:10.1016/S0893-6080(98)00010-0.

[53] Fawcett T. An introduction to ROC analysis. Pattern Recognit Lett. 2006;27(8):861–874. doi:10.1016/j.patrec.2005.10.010.

[54] Yousefabadi H, Mehrmohamadi M. Bioactivity-driven prediction of antibacterial synergy using machine learning models. bioRxiv preprint. doi:10.64898/2025.12.14.694088.

[55] Jolliffe IT, Cadima J. Principal component analysis: a review and recent developments. Philosophical Transactions of the Royal Society A: Mathematical, Physical and Engineering Sciences. 2016;374(2065):20150202. doi:10.1098/rsta.2015.0202.

[56] Chan DI, Prenner EJ, Vogel HJ. Tryptophan- and arginine-rich antimicrobial peptides: structures and mechanisms of action. Biochimica et Biophysica Acta (BBA) - Biomembranes. 2006;1758(9):1184–1202. doi:10.1016/j.bbamem.2006.04.006.

[57] Hilpert K, Volkmer-Engert R, Walter T, Hancock REW. High-throughput generation of small antibacterial peptides with improved activity. Nature Biotechnology. 2005;23(8):1008–1012. doi:10.1038/nbt1113.

[58] Hilpert K, Elliott MR, Volkmer-Engert R, Henklein P, Donini O, Zhou Q, Winkler DFH, Hancock REW. Sequence requirements and an optimization strategy for short antimicrobial peptides. Chemistry & Biology. 2006;13(10):1101–1107. doi:10.1016/j.chembiol.2006.08.014.

[59] Remington JM, Liao C, Sharafi M, Ste Marie EJ, Ferrell JB, Hondal RJ, Wargo MJ, Schneebeli ST, Li J. Aggregation state of synergistic antimicrobial peptides. The Journal of Physical Chemistry Letters. 2020;11(21):9501–9506. doi:10.1021/acs.jpclett.0c02094.

[60] Sánchez-Gómez S, Japelj B, Jerala R, Moriyón I, Fernández Alonso M, Leiva J, Blondelle SE, Andrä J, Brandenburg K, Lohner K, Martínez de Tejada G. Structural features governing the activity of lactoferricin-derived peptides that act in synergy with antibiotics against *Pseudomonas aeruginosa* in vitro and in vivo. Antimicrobial Agents and Chemotherapy. 2011;55(1):218–228. doi:10.1128/AAC.00904-10.

[61] Li XZ, Livermore DM, Nikaido H. Role of efflux pump(s) in intrinsic resistance of *Pseudomonas aeruginosa*: resistance to tetracycline, chloramphenicol, and norfloxacin. Antimicrobial Agents and Chemotherapy. 1994;38(8):1732–1741. doi:10.1128/AAC.38.8.1732.

[62] Li XZ, Nikaido H, Poole K. Role of MexA-MexB-OprM in antibiotic efflux in *Pseudomonas aeruginosa*. Antimicrobial Agents and Chemotherapy. 1995;39(9):1948–1953. doi:10.1128/AAC.39.9.1948.

[63] Masuda N, Sakagawa E, Ohya S, Gotoh N, Tsujimoto H, Nishino T. Contribution of the MexX-MexY-OprM efflux system to intrinsic resistance in Pseudomonas aeruginosa. Antimicrobial Agents and Chemotherapy. 2000;44(9):2242–2246. doi:10.1128/AAC.44.9.2242-2246.2000.

[64] Wesseling CMJ, Martin NI. Synergy by perturbing the Gram-negative outer membrane: opening the door for Gram-positive specific antibiotics. ACS Infectious Diseases. 2022;8(9):1731–1757. doi:10.1021/acsinfecdis.2c00193.

[65] Dhiman S, Ramirez D, Arora R, Arthur G, Schweizer F. Enhancing outer membrane permeability of tetracycline antibiotics in *P. aeruginosa* using TOB-CIP conjugates. RSC Medicinal Chemistry. 2024;15:3133–3146. doi:10.1039/D4MD00329B.

[66] de Sousa T, Silva C, Alves O, Costa E, Igrejas G, Poeta P, Hébraud M. Determination of antimicrobial resistance and the impact of imipenem + cilastatin synergy with tetracycline in *Pseudomonas aeruginosa* isolates from sepsis. Microorganisms. 2023;11(11):2687. doi:10.3390/microorganisms11112687.

[67] Bordes A, Usunier N, Garcia-Duran A, Weston J, Yakhnenko O. Translating embeddings for modeling multi-relational data. In: Advances in Neural Information Processing Systems 26. 2013;2787–2795.

[68] Sun Z, Deng ZH, Nie JY, Tang J. RotatE: Knowledge graph embedding by relational rotation in complex space. International Conference on Learning Representations. 2019.

[69] Kipf TN, Welling M. Semi-supervised classification with graph convolutional networks. International Conference on Learning Representations. 2017.

[70] Klicpera J, Bojchevski A, Günnemann S. Predict then propagate: Graph neural networks meet personalized PageRank. In: International Conference on Learning Representations (ICLR); 2019.

[71] Velickovic P, Cucurull G, Casanova A, Romero A, Liò P, Bengio Y. Graph attention networks. In: International Conference on Learning Representations (ICLR); 2018.

[72] Xu K, Hu W, Leskovec J, Jegelka S. How powerful are graph neural networks? In: International Conference on Learning Representations (ICLR); 2019.

[73] Hu W, Liu B, Gomes J, Zitnik M, Liang P, Pande VS, Leskovec J. Strategies for pre-training graph neural networks. In: International Conference on Learning Representations (ICLR); 2020.

[74] Shi Y, Huang Z, Feng S, Zhong H, Wang W, Sun Y. Masked label prediction: unified message passing model for semi-supervised classification. In: Proceedings of the Thirtieth International Joint Conference on Artificial Intelligence (IJCAI-21). 2021;1548–1554. doi:10.24963/ijcai.2021/214.

[75] Sánchez-Gómez S, Ferrer-Espada R, Stewart PS, Pitts B, Lohner K, Martínez de Tejada G. Antimicrobial activity of synthetic cationic peptides and lipopeptides derived from human lactoferricin against *Pseudomonas aeruginosa* planktonic cultures and biofilms. BMC Microbiology. 2015;15:137. doi:10.1186/s12866-015-0473-x.

[76] Ruden S, Rieder A, Chis Ster I, Schwartz T, Mikut R, Hilpert K. Synergy pattern of short cationic antimicrobial peptides against multidrug-resistant *Pseudomonas aeruginosa*. Frontiers in Microbiology. 2019;10:2740. doi:10.3389/fmicb.2019.02740.

[77] Jahangiri A, Neshani A, Mirhosseini SA, Ghazvini K, Zare H, Sedighian H. Synergistic effect of two antimicrobial peptides, Nisin and P10 with conventional antibiotics against extensively drug-resistant *Acinetobacter baumannii* and colistin-resistant *Pseudomonas aeruginosa* isolates. Microbial Pathogenesis. 2021;150:104700. doi:10.1016/j.micpath.2020.104700.

[78] Olsson A, Wistrand-Yuen P, Nielsen EI, Friberg LE, Sandegren L, Lagerbäck P, Tängdén T. Efficacy of antibiotic combinations against multidrug-resistant *Pseudomonas aeruginosa* in automated time-lapse microscopy and static time-kill experiments. Antimicrobial Agents and Chemotherapy. 2020;64(6):e02111–19. doi:10.1128/AAC.02111-19.

[79] Ly NS, Bulman ZP, Bulitta JB, Baron C, Rao GG, Holden PN, Li J, Sutton MD, Tsuji BT. Optimization of polymyxin B in combination with doripenem to combat mutator *Pseudomonas aeruginosa*. Antimicrobial Agents and Chemotherapy. 2016;60(5):2870–2880. doi:10.1128/AAC.02377-15.

[80] Domalaon R, Sanchak Y, Koskei LC, Lyu Y, Zhanel GG, Arthur G, Schweizer F. Short proline-rich lipopeptide potentiates minocycline and rifampin against multidrug- and extensively drug-resistant *Pseudomonas aeruginosa*. Antimicrobial Agents and Chemotherapy. 2018;62(4):e02374–17. doi:10.1128/AAC.02374-17.

[81] Lister PD, Wolter DJ, Hanson ND. Antibacterial-resistant *Pseudomonas aeruginosa*: clinical impact and complex regulation of chromosomally encoded resistance mechanisms. Clinical Microbiology Reviews. 2009;22(4):582–610. doi:10.1128/CMR.00040-09.

[82] Breiman L. Random forests. Mach Learn. 2001;45(1):5–32. doi:10.1023/A:1010933404324.

[83] Cox DR. The regression analysis of binary sequences. J R Stat Soc Series B Stat Methodol. 1958;20(2):215–242. doi:10.1111/j.2517-6161.1958.tb00292.x.

